# Application of TurboID-mediated proximity labeling for mapping a GSK3 kinase signaling network in Arabidopsis

**DOI:** 10.1101/636324

**Authors:** Tae-Wuk Kim, Chan Ho Park, Chuan-Chih Hsu, Jia-Ying Zhu, Yuchun Hsiao, Tess Branon, Shou-Ling Xu, Alice Y Ting, Zhi-Yong Wang

## Abstract

Transient protein-protein interactions (PPIs), such as those between posttranslational modifying enzymes and their substrates, play key roles in cellular regulation, but are difficult to identify. Here we demonstrate the application of enzyme-catalyzed proximity labeling (PL), using the engineered promiscuous biotin ligase TurboID, as a sensitive method for characterizing PPIs in signaling networks. We show that TurboID fused with the GSK3-like kinase BIN2 or a PP2A phosphatase biotinylates their known substrate, the BZR1 transcription factor, with high specificity and efficiency. We optimized the protocol of biotin labeling and affinity purification in transgenic Arabidopsis expressing a BIN2-TurboID fusion protein. Subsequent quantitative mass spectrometry (MS) analysis identified about three hundred proteins biotinylated by BIN2-TurboID more efficiently than the YFP-TurboID control. These include a significant subset of previously proven BIN2 interactors and a large number of new BIN2-proximal proteins that uncover a broad BIN2 signaling network. Our study illustrates that PL-MS using TurboID is a powerful tool for mapping signaling networks, and reveals broad roles of BIN2 kinase in cellular signaling and regulation in plants.

**Impact Statement:** TurboID-mediated proximity labeling is a powerful tool for protein interactomics in plants.

## Introduction

Protein-protein interactions (PPIs) organize proteins into cellular structures, metabolic complexes, and regulatory pathways, and are therefore of fundamental importance for understanding both cellular processes and developmental programs (Gingras et al., 2019, Bontinck et al., 2018). Interactions between different proteins have different affinity and kinetics to satisfy the dynamic needs of cellular machineries and regulatory pathways. For example, subunits of enzymatic or structural complexes tend to form stable interactions, whereas interactions between components of signaling pathways tend to be transient and dynamic. Elucidation of cellular PPIs, particularly transient PPIs in signaling networks, remains a major challenge in biology (Gingras et al., 2019).

The most commonly-used methods for identification of interacting proteins include the yeast two-hybrid (Y2H) screen and immunoprecipitation (IP) followed by mass spectrometry (MS). Y2H screen identifies pairwise interactions in yeast, and has been used traditionally for identifying interactors of individual proteins and more recently for generating large scale interactome networks (Arabidopsis Interactome Mapping, 2011, Jones et al., 2014, Trigg et al., 2017). IP-MS captures endogenous proteins that interact with bait protein *in vivo*, including proteins bound directly and indirectly in a protein complex (Bontinck et al., 2018). While Y2H and IP-MS are widely used, both have major problems of high rates of false positive and false negative artifacts for various reasons (Braun et al., 2013, Bontinck et al., 2018, Gingras et al., 2019). Interactions detected by Y2H usually need verification by *in vivo* assays such as co-immunoprecipitation. In IP-MS, false positives can occur when proteins of separate cellular compartments are mixed in the extract or, more often, when non-specific bindings are not removed by washing or quantitative comparison to proper negative controls. IP-MS is particularly problematic for identifying transient PPIs, which is often the case between signaling proteins, as such interactions tend to be lost during the procedures of extraction, incubation, and washing of the beads. The problem is especially challenging for proteins that require harsh extraction conditions, such as membrane proteins requiring detergent to extract (Bontinck et al., 2018, Gingras et al., 2019).

*In vivo* affinity tagging by proximity-based biotinylation, an approach named proximity labeling (PL), has emerged as an alternative to IP for proteomic analysis of PPIs in animal systems (Roux et al., 2012, Lobingier et al., 2017, Han et al., 2017, Han et al., 2018, Li et al., 2017, Lönn and Landegren, 2017, Gingras et al., 2019). Enzymes used for PL include an engineered soybean ascorbate peroxidase (named APEX2) (Rhee et al., 2013), and a promiscuous mutant of the *Escherichia coli* biotin ligase BirA (named BioID). These enzymes are expressed in cells as fusions with a protein of interest, and catalyze biotinylation of proximal and interacting proteins (Roux et al., 2012, Rhee et al., 2013). The biotin-labeled proteins can be affinity purified by streptavidin pulldown and subsequently identified by MS (Roux et al., 2012). PL-MS has been used extensively in animal systems to map protein interaction networks, organelle proteomes, and spatiotemporally resolved protein interaction networks in living cells (Uezu et al., 2016, Kim et al., 2014, Roux et al., 2012, Lobingier et al., 2017, Gingras et al., 2019). However, application of PL-MS in plants has yielded very limited success (Conlan et al., 2018, Khan et al., 2018, Lin et al., 2017, Bontinck et al., 2018), potentially due to the low catalytic activity of BioID and, for APEX2, high background due to endogenous peroxidase activity.

Recently, Branon *et al*. dramatically improved the tagging kinetics of BioID by directed evolution of BirA (Branon et al., 2018). The new promiscuous BirA mutant, dubbed TurboID, catalyzes PL at much greater efficiency than BioID (Branon et al., 2018). After ten minutes of biotin feeding, TurboID achieved a level of biotinylation that required 18 hours of biotin feeding with BioID (Branon et al., 2018). Targeting TurboID to subcellular compartments such as mitochondria, ER, and nucleus in Drosophila successfully identified the proteomes of these organelles (Branon et al., 2018). The effectiveness of TurboID in mapping signaling networks remains to be explored, and the application of TurboID in plants has not been reported.

Brassinosteroid (BR) is a major growth-promoting hormone, and it acts through one of the best-characterized receptor kinase-mediated signal transduction pathway in plants. BR binding to the BRI1 receptor kinase triggers a cascade of signaling events that include phosphorylation of members of the BSK family kinases and the BSU family phosphatases, BSU1-mediated dephosphorylation and inactivation of GSK3-like kinases including BIN2, and PP2A-mediated dephosphorylation of the BZR1 family transcription factors (Tang et al., 2008, Kim et al., 2009, Kim and Wang, 2010). When BR levels are low, BIN2 phosphorylates BZR1 to inhibit its nuclear localization and modulate BR-responsive gene expression. BIN2 has also been reported to phosphorylate additional transcription factors and kinases to modulate other signaling and developmental pathways (Vert et al., 2008, Kim et al., 2012, Cai et al., 2014, Cho et al., 2014, Kondo et al., 2014). A complete understanding of the BIN2 signaling network would require identification of all the BIN2 substrates and regulators *in vivo* (Youn and Kim, 2015).

Considering the high catalytic activity of TurboID, we wondered if it could be an efficient tool for capturing transient and dynamic interactors of a bait protein like BIN2 in plants. We reasoned that a fusion between TurboID and a kinase or phosphatase will biotinylate the substrates of the enzyme during the process of them being phosphorylated or de-phosphorylated, allowing their identification by PL-MS. Our results show that BZR1 can be specifically biotinylated by TurboID fusions with BIN2 or the phosphatase PP2AB’α subunit, which are known to interact with BZR1 (He et al., 2002, Tang et al., 2011). After optimizing the conditions for biotin feeding, protein extraction and streptavidin pulldown, we performed PL-MS of BIN2-TurboID and successfully identified a large number of BIN2-interacting and proximal proteins, which include a significant fraction of the BIN2 interactors reported previously with *in vivo* validation, demonstrating a high sensitivity of TurboID-mediated proximity labeling in plants. The results further provide evidence for a very large BIN2-mediated signaling network. Our study demonstrates the application of TurboID as a powerful tool for mapping signaling networks, and establishes the framework of a large GSK3/BIN2-signaling network in Arabidopsis.

## Results

To investigate whether TurboID is an effective tool for PL in plant cells (Figure 1A), we generated a gateway-compatible binary vector to express proteins fused to TurboID (Figure 1B). We cloned the full-length coding sequence of BIN2 into the vector to express a fusion protein containing BIN2, YFP (yellow florescent protein) and TurboID (BIN2-YFP-TbID) from the constitutive 35S promoter. To test whether BIN2-YFP-TbID can biotinylate BZR1 in plant cells, we co-expressed it with a BZR1 fused with 4xMyc-6His tag (BZR1-MH) transiently in *Nicotiana benthamiana* leaves. As a control, we also co-expressed BZR1-MH with a YFP-YFP-TbID fusion protein, using the constitutive 35S promoter. After pull down by Streptavidin-agarose from the protein extracts of *Nicotiana benthamiana* leaves, immunoblotting with anti-Myc antibody detected BZR1-MH in the sample that co-expressed BIN2-YFP-TbID but not in the YFP-YFP-TbID control (Figure 1C), indicating that BIN2 interaction with BZR1 can be detected specifically by TurboID-based PL in plant cells.

**Figure 1.**
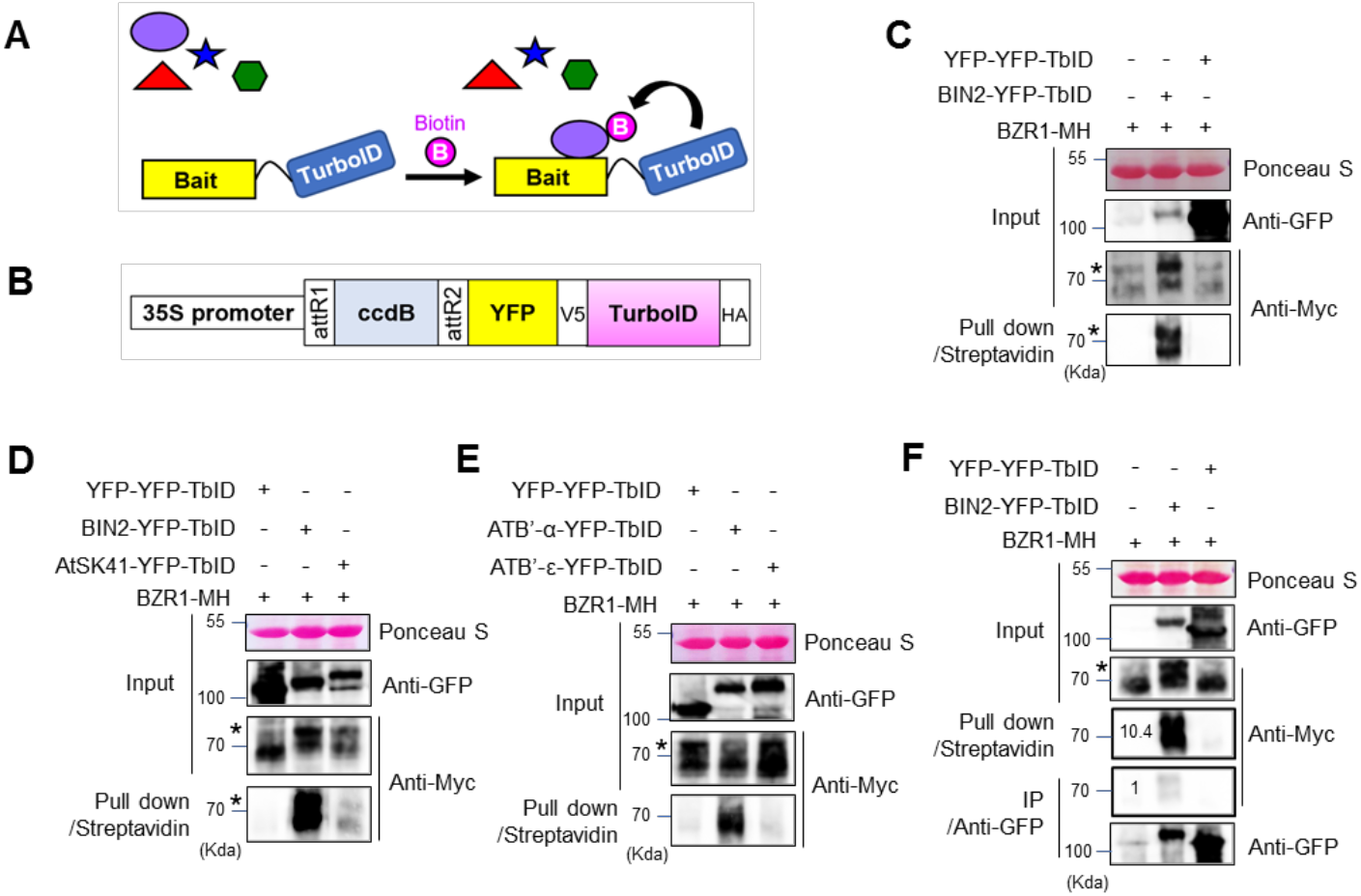
Investigation of TurboID-based biotin labeling for detecting interactions between brassinosteroid signaling components. (A) Schematic diagram of proximity-dependent biotin labeling of interacting proteins. (B) Diagram of the Gateway-compatible binary vector used in this study to express YFP-tagged TurboID (TbID) fusion protein (*ccdB* to be replace with coding sequence of interest, such as BIN2). (C) TbID-based detection of the interaction between BIN2 and BZR1. Agrobacteria carrying *35S:YFP-YFP-TbID* or *35S:BIN2-YFP-TbID* was co-infiltrated with that carrying *35S:BZR1-4Myc-6His* (MH) into *N. benthamiana* leaves. BZR1-MH pulled down by streptavidin agarose was detected by anti-Myc antibody. (D) Comparison of BZR1 biotinylation by BIN2 and AtSK41. BZR1-MH was co-expressed with YFP-YFP-TbID, BIN2-YFP-TbID, or AtSK41-YFP-TbID, and then analyzed as in panel C. (E) Comparison of BZR1 biotinylation by PP2AB’α and PP2AB’ε. BZR1-MH was co-expressed with YFP-YFP-TbID, PP2AB’α (ATB’-α)-YFP-TbID or PP2AB’ε (ATB’-ε)-YFP-TbID, and analyzed as in panel C. (F) Proximity labeling is more efficient than Co-IP for detecting the *in vivo* interaction between BIN2 and BZR1. BZR1-MH co-expressed in *N. benthamiana* leaves with BIN2-YFP-TbID, or with YFP-YFP-TbID as a negative control, was pulled down with streptavidin-agarose or co-immunoprecipitated by anti-GFP antibody from aliquots of the same protein extracts, and then immunoblotted using anti-Myc antibody. Numbers indicate relative signal level of BZR1-MH normalized to the signal obtained by Co-IP. (C-F) Asterisks indicate phosphorylated BZR1.

We further examined the specificity of TurboID for biotinylation of interacting proteins. Previous studies demonstrated that BZR1 interacts with six GSK3 kinases including BIN2 but not with the other three GSK3s such as AtSK41 in Y2H assays (Kim et al., 2009). Consistently, BZR1-MH biotinylation by BIN2-YFP-TbID was much stronger than that by AtSK41-YFP-TbID (Figure 1D). Previous studies also showed that BZR1 binds to specific PP2A regulatory subunits such as PP2AB’α, but not to PP2AB’ε (Tang et al., 2011). Again, biotinylation of BZR1-MH was detected when co-expressed with PP2AB’α-YFP-TbID but not with PP2AB’ε-YFP-TbID (Figure 1E). Our results indicate that PL using TurboID has a relatively high level of specificity for true interacting proteins.

To compare the efficiency of PL with co-immunoprecipitation (co-IP), protein extracts obtained from leaves co-expressing BZR1-MH and BIN2-YFP-TbID or leaves co-expressing BZR1-MH and YFP-YFP-TbID were evenly divided into two aliquots. Each aliquot was pulled down with streptavidin-agarose beads or with anti-GFP antibody immobilized on Protein A-agarose beads. The streptavidin beads pulled down about 10 times more BZR1-MH than did the anti-GFP antibody beads (Figure 1F), indicating that TurboID-based PL is more efficient than co-IP for detecting PPI in plant cells.

To further identify BIN2-interacting proteins using PL-MS, we generated transgenic Arabidopsis plants expressing BIN2-YFP-TbID or YFP-YFP-TbID as a control. The plants expressing YFP-YFP-TbID displayed normal plant growth (Figure 2A). Notably, some of the plants expressing *BIN2-YFP-TbID* displayed typical dwarf phenotypes, as observed in BIN2-overexpressing or *bin2-1* mutant plants (Li and Nam, 2002), suggesting that BIN2-YFP-TbID is functional in Arabidopsis (Figure 2A–2B and Figure 2-figure supplement 1). In addition, confocal microscopic analysis confirmed that YFP-YFP-TbID and BIN2-YFP-TbID are similarly localized in the cytoplasm and the nucleus consistent with previously reported BIN2 localization (Figure 2C–2D) (Kim et al., 2009).

**Figure 2.**
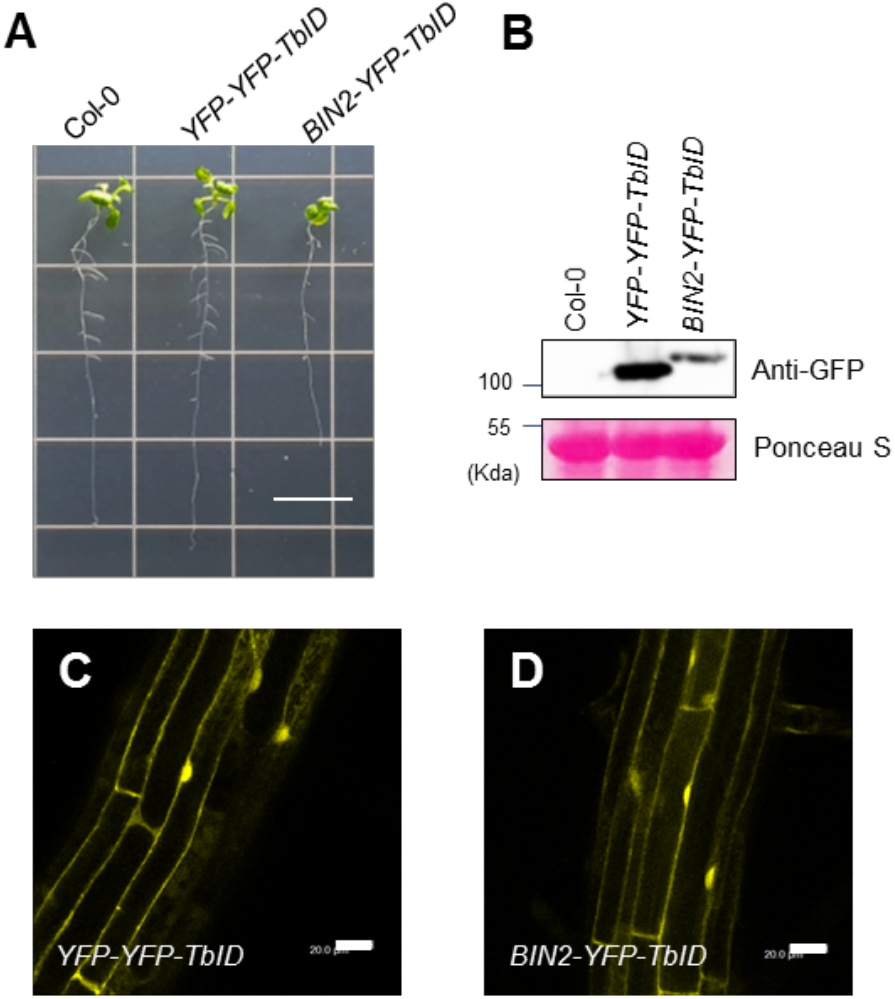
Phenotype and subcellular localization of *YFP-YFP-TbID* and *BIN2-YFP-TblD*. (**A**) Phenotype of the wild-type (Col-0) and transgenic Arabidopsis overexpressing *YFP-YFP-TbID* or *BIN2-YFP-TbID*. Seedlings were grown for 14 days on agar medium. Scale bar indicates 1 cm. (**B**) Immunoblot analysis for protein expression in the seedlings shown in (**A**). (**C-D**) Subcellular localization of YFP-YFP-TbID (**C**) and BIN2-YFP-TbID (**D**) in Arabidopsis. Roots of 8-day-old seedlings grown MS medium in the light were observed by confocal microscope. Bars = 20 μm.

We tested the effects of different biotin treatment conditions on the efficiencies of biotinylation and streptavidin pulldown. We found that biotin treatment increased the biotinylation of proteins in the *BIN2-YFP-TbID* Arabidopsis seedlings in a concentration- and time-dependent manner up to 250 μM biotin for 1 hr (Figure 3A), or six hours of treatment with 50 μM biotin (Figure 3B). We then tested the effects of biotin concentration on streptavidin pulldown. We found that 5 μM biotin treatment significantly increased the amount of BZR1-MH pulled down by streptavidin-agarose (Figure 3C). However, unexpectedly, treatments with higher concentration (50 and 500 μM) of biotin greatly reduced the efficiency of pulldown. Immunoblotting using anti-Myc antibody showed that the level of BZR1-MH protein was not changed, and its biotinylation was increased upon biotin treatment at the higher concentrations (Figure 3C). We suspected that the high concentrations of exogenous biotin enhance PL but interfere with the pulldown by streptavidin if not removed from the extracts. We investigated whether removal of free biotin from the extracts can improve the efficiency of pulldown by streptavidin. An aliquot of the extract of *BIN2-YFP-TbID* seedlings treated with 50 μM biotin was desalted using a Sephadex G-25 desalting column. The removal of free biotin by desalting step greatly increased the pulldown efficiency of biotinylated proteins, whereas no appreciable amount of biotinylated proteins was pulled down without desalting treatment (Figure 3D).

**Figure 3.**
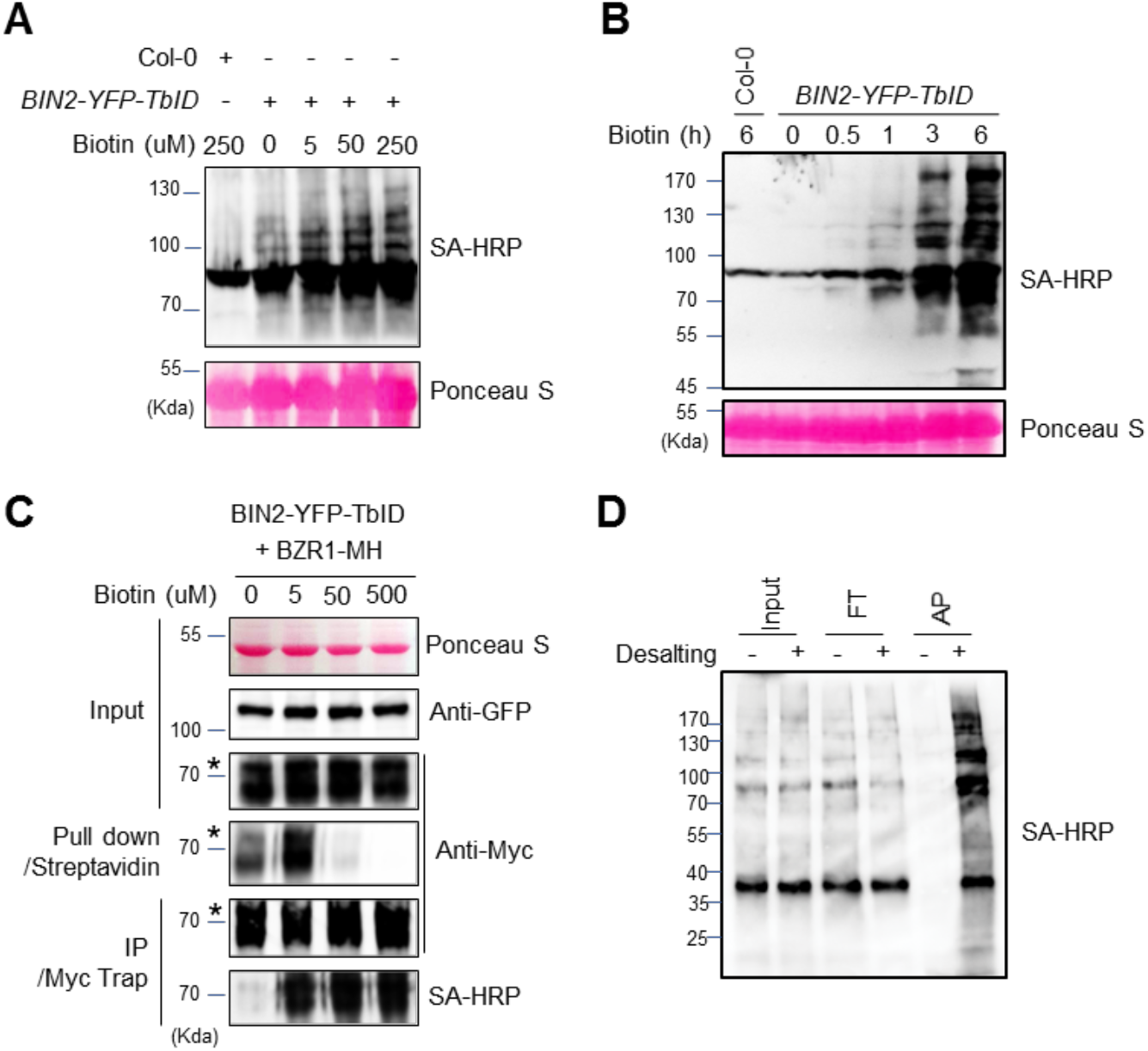
The effect of biotin concentration and incubation time on the efficiency of biotin labeling and affinity purification. (**A**) The effect of biotin concentration on TbID-mediated biotinylation. *BIN2-YFP-TbID* seedlings were treated by the indicated concentrations of biotin for 1 h. Total proteins were gel-blotted using biotinylation by streptavidin-HRP (SA-HRP) to detect protein biotinylation. (**B**) *BIN2-YFP-TbID* transgenic seedlings were grown on medium for 14 days, and then treated with 50 μM biotin for the indicated time, and analyzed as in panel A. (**C**) Effects of biotin concentration on streptavidin pulldown of biotinylated proteins. *N. benthamiana* leaves co-expressing BIN2-YFP-TbID and BZR1-MH were treated by biotin of the indicated concentrations for 30 min. The proteins pulled down with streptavidin-agarose or immunoprecipitated by anti-Myc antibody, were gel-blotted using anti-Myc antibody and SA-HRP. (**D**) The effect of desalting on affinity purification of biotinylated proteins. *BIN2-YFP-TbID* seedlings were treated with 50 μM biotin for 3 h and aliquots of the protein extract were treated with (+) or without (−) desalting step. Input (1:300), flow through (FT, 1:300) and eluate after streptavidin affinity purification (AP, 1:10) were gel blotted using SA-HRP.

We further tested the effects of different concentrations of biotin on the identification of biotinylated proteins in PL-MS. We treated the *BIN2-YFP-TbID* seedlings with mock solution, or 5, 50, or 500 μM of biotin for one hour. Total proteins were extracted from *BIN2-YFP-TbID* plants, precipitated by methanol-chloroform, and digested by trypsin. The peptides were incubated with streptavidin beads and the bound peptides were eluted and analyzed by LC-MS/MS. The number of biotinylated peptides increased with increasing concentrations of biotin: from 365 biotinylated peptides of 268 proteins detected in the mock treated samples to 1485 peptides of 620 proteins detected in the sample treated with 500 μM of biotin (Figure 4A). Most of the peptides detected at no or low concentrations of biotin (304 biotinylated peptides) were also detected when higher concentrations of biotin were used (Figure 4B, Figure 4-figure supplement 1). When the signal intensities were analyzed, we found that the peptides detected with endogenous biotin showed a saturation of biotinylation at 5 μM biotin and their biotinylation did not further increase at higher biotin concentrations (Figure 4C), whereas the peptides detected at 5 μM or 50 μM biotin showed further increase of biotinylation at higher biotin concentrations (Figure 4D). These observations suggest that the dosage of biotin treatment has major effects on the sensitivity and specificity, and a proper negative control is essential for distinguishing real from false interactors.

**Figure 4.**
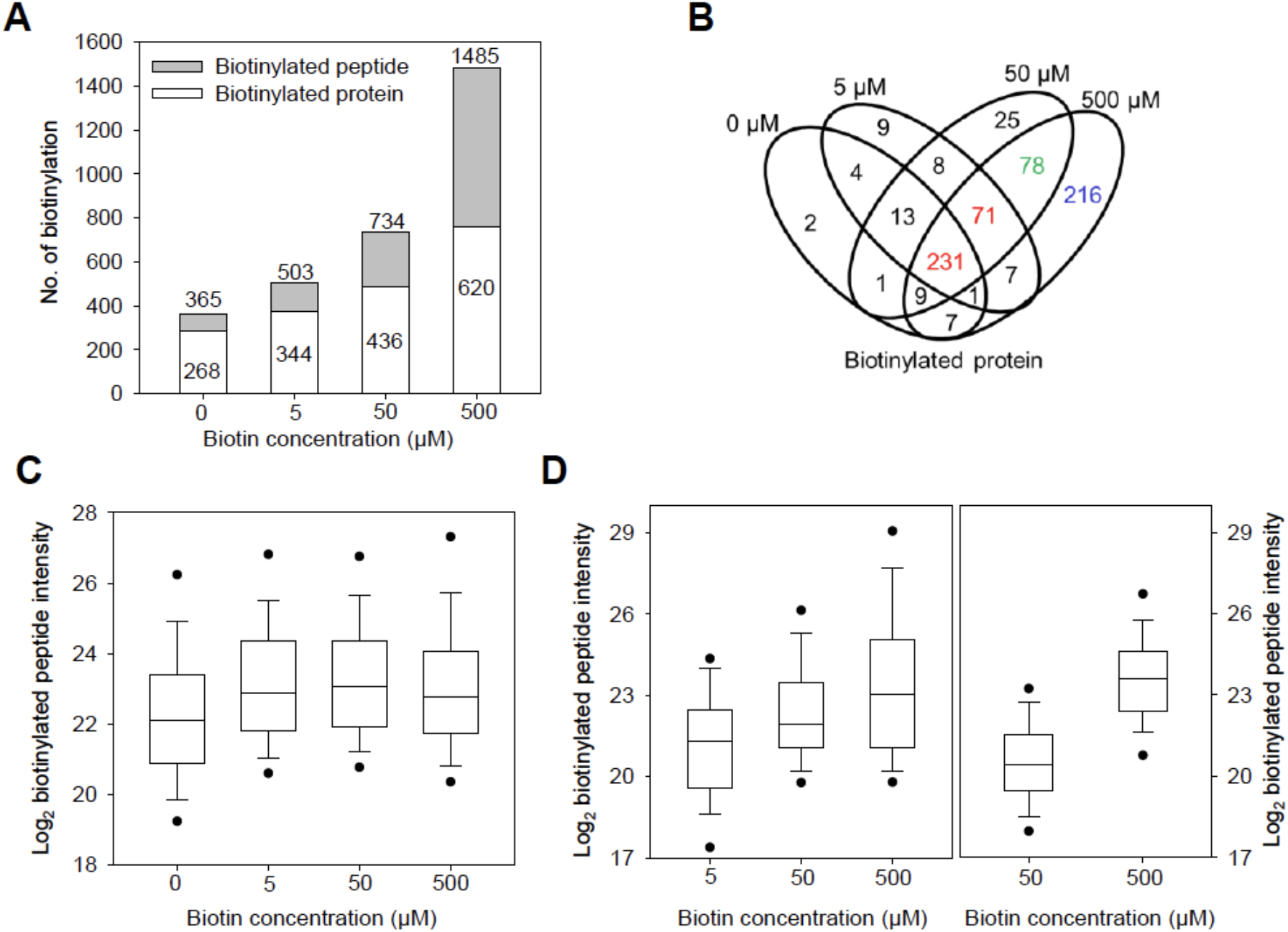
The effect of biotin concentration on the identification of biotinylated proteins using mass spectrometry. (**A**) The number of identified biotinylated peptides and proteins are increased with increased concentration of biotin. (**B**) The overlap of identified biotinylated proteins between 1 h of 0, 5, 50, and 500 μM biotin-treated samples. The biotinylated proteins identified in all four, three (5~500 μM), two (50~500 μM), and only 500 μM concentrations are 231 proteins, 71 proteins, 78 proteins, and 216 proteins, respectively. (**C**) Comparison of the signal intensities of biotinylated peptides identified from all four biotin concentrations (0~500 μM). The overall intensity of these biotinylated peptides is saturated at 5 μM of biotin concentration. (**D**) Comparison of the intensities of biotinylated peptides identified from three of biotin concentrations (left panel) and two of biotin concentrations (right panel). The intensity of biotinylated peptide is increased with higher biotin concentration.

With the method optimized, we tried to identify the BIN2 interactome using transgenic Arabidopsis expressing BIN2-YFP-TbID with those expressing YFP-YFP-TbID as negative control. Considering the importance of quantitative comparison to the negative control, we chose to use metabolic stable isotope labeling mass spectrometry (mSIL-MS) to quantitatively analyze the proteins biotinylated in *BIN2-YFP-TbID* relative to the *YFP-YFP-TbID* control. We grew the *BIN2-YFP-TbID* and *YFP-YFP-TbID* seedlings on Murashige and Skoog medium containing either an ^14^N or ^15^N nitrogen source for 16 days, then treated the seedlings with 50 μM biotin for 3 hours. Free biotin was removed from protein extracts by desalting column, and biotinylated proteins were affinity purified using streptavidin beads (Figure 5-figure supplement 1). The proteins of the ^14^N-labeled *YFP-YFP-TbID* sample were mixed with those of ^15^N-labeled *BIN2-YFP-TbID* seedlings. In a biological repeat, ^15^N-labeled *YFP-YFP-TbID* and ^14^N-labeled *BIN2-YFP-TbID* were mixed. The proteins were separated in SDS-PAGE and in-gel digested by trypsin. The peptides were analyzed by liquid chromatography (LC)-MS/MS, and the ratios between the isotope-coded peptides were quantified as described in Material and Methods (Figure 5A). The results show that the bait protein BIN2 was only detectable in the *BIN2-YFP-TbID* samples as expected (Figure 5B). While most of the detected proteins, such as MLP34, showed no significant difference between *YFP-YFP-TbID* and *BIN2-YFP-TbID* (Figure 5B), a total of 55 proteins were consistently enriched over 2-fold in *BIN2-YFP-TbID* relative to the *YFP-YFP-TbID* control in both replicate experiments (Figure 5C, Figure 5-Source data 1). These include known BIN2 interacting protein families such as BSK (BSK3) and BSUs (BSL2/BSL3) (Figure 5B and 5C). In addition, the quantitative data and mass spectra showed dramatic enrichment of many proteins previously unknown for interaction with BIN2, such as the auxin transporter PIN3, NONPHOTOTROPIC HYPOCOTYL3 (NPH3), and two receptor kinases HERK1 and FER (Figure 5B and 5C).

**Figure 5.**
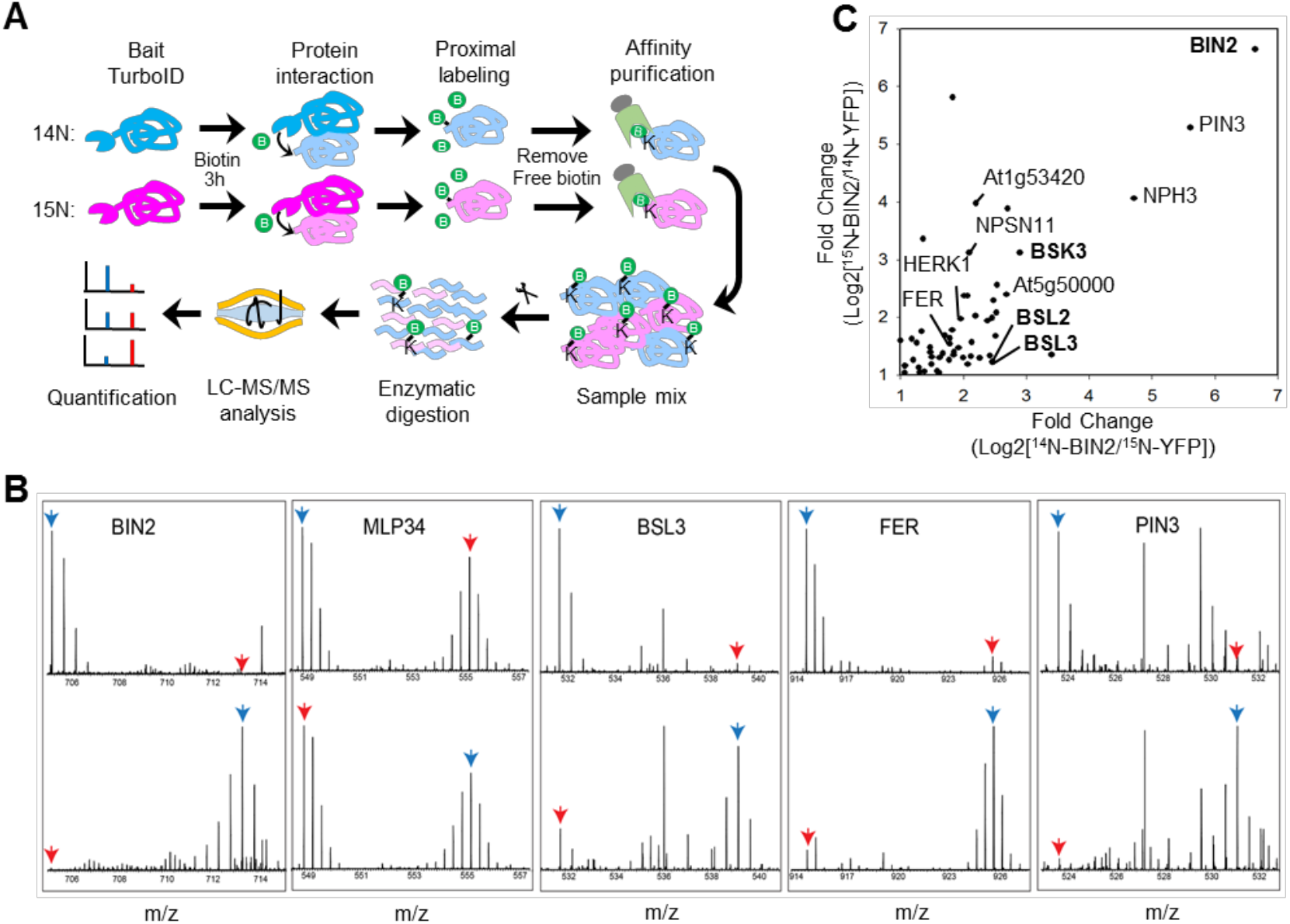
TbID-based identification of BIN2-interacting proteins in Arabidopsis. (**A**) Schematic diagram of biotinylated-protein affinity purification-mass spectrometry analysis. (**B**) MS1 peptide spectra show the enrichment of BIN2, BSL3, FERONIA (FER) and PIN3 in PL-MS of BIN2-YFP-TbID vs. YFP-YFP-TbID control, and no enrichment of MLP34, a non-specific interactor. Top panel: ^14^N-BIN2-YFP-TbID, ^15^N-YFP-YFP-TbID; Bottom panel: ^14^N-YFP-YFP-TbID, ^15^N-BIN2-YFP-TbID. Blue arrows point to BIN2 samples, red arrows point to control samples. (**C**) Scatter plot of log2 folds enrichment (signal ration between BIN2-YFP-TbID and YFP-YFP-TbID) of proteins from two replicate experiments where isotopes were switched. Bold letter indicates previously reported BIN2 interactors

The mass spectrometry analysis detected biotinylation in only a very small fraction of the identified proteins, suggesting that a large number of non-biotinylated background proteins were pulled down in the above experiment. We thus tested whether streptavidin pulldown after trypsin digestion could identify more biotinylated peptides of real interactors. Total proteins were extracted from *BIN2-YFP-TbID* and *YFP-YFP-TbID* plants treated with 50 μM biotin for 3 hrs, precipitated by methanol-chloroform, and digested by trypsin. The peptides were divided into three replicates and then were incubated with streptavidin beads and the bound peptides were eluted and analyzed by LC-MS/MS. We identified 12,441 and 13,805 unique peptides, of which 463 and 1,073 were biotinylated from the *YFP-YFP-TbID* and *BIN2-YFP-TbID* plants, respectively. Among these, 375 biotinylated peptides from 280 proteins (in addition to BIN2) were identified in all three replicates of *BIN2-YFP-TbID* but not in *YFP-YFP-TbID*, and they are considered BIN2-proximal proteins (Figure 5-Source data 2). These proteins include additional known BIN2 interactors, such as BZR1, BES1, OPS, YDA, and ARF2, which were not detected in the protein pulldown experiment. Of the 55 proteins detected in the protein pulldown experiments, 25 were detected in the peptide purification experiments, and 24 of these 25 showed over 5-fold enrichment in BIN2-YFP-TbID compared to the YFP-YFP-TbID control (Figure 5-Source data 1). Together these experiments identified 310 BIN2-proximal proteins, which include eight previously reported BIN2-interactors.

## Discussion

While PPIs are of fundamental importance for all cellular and developmental processes, mapping PPIs network is technically challenging, particularly for transient and dynamic interactions, which often play important regulatory roles (Bontinck et al., 2018). In particular, the interactions between posttranslational modifying enzymes, such as kinases and phosphatases, and their substrate proteins tend to be transient and thus difficult to capture in traditional IP-MS approaches. Here we show that the super-active biotin ligase TurboID fused with a kinase or phosphatase can biotinylate their substrate proteins, demonstrating the application of TurboID as an effective tool for mapping signaling networks.

IP-MS has been the common approach to mapping *in vivo* PPIs. A major challenge in IP-MS experiment is to maintain PPIs while removing non-specific background proteins. In order to minimize non-specific background, extensive washes using harsh conditions are often used, and even tandem affinity purification have been developed (Trinkle-Mulcahy, 2019). These kinds of efforts to reduce background lead to loss of some real interactors, especially dynamic interactors. Accordingly, mild wash conditions combined with quantitative MS analysis of enrichment has been used (Wei et al., 2016; Ni et al., 2014). However, identification of real interactors can be compromised by the large amount of non-specific proteins that overwhelm the MS analysis (Wei et al., 2016, Bu et al., 2017). While most cellular proteins are phosphorylated, only a small number of them have been shown to interact with any kinase using current technologies. Apparently, more sensitive methods are required to map the kinase-substrate interactions. Our study demonstrates that TurboID-mediated PL-MS is a potential solution for this challenging problem in biology.

PL has several advantages over the traditional AP approaches for identifying *in vivo* PPIs, including high-affinity capture of *in vivo* interactions with minimum effect of protein extraction conditions (Gingras et al., 2019, Han et al., 2018). One major advantage is detecting dynamic and transient interactions, where PL benefits from a signal amplification or accumulation effect. The amount of an interacting protein captured in AP experiment is dependent on the concentrations of the interactors, the binding affinity and off rate of the interaction, and time of incubation and washing. Dynamic interactions that rapidly cycle the interacting partners tend to be lost during the washing steps. Some interactions maybe disrupted by the extraction and washing conditions that are different from the cellular environment, such as detergent used to extract membrane proteins. In PL, processive biotinylation of dynamic interactors over the time period of biotin treatment (and with endogenous biotin over the growth period) can accumulate a much larger pool of biotin-tagged interactors than the amount bound to the bait at a steady state. Such accumulated biotin-tagged proteins can be purified with high affinity of streptavidin and identified by MS. This advantage of PL for identifying dynamic PPIs has not been explored for mapping post-translational modification and signaling networks, where dynamic PPIs are prominent and important (Gingras et al., 2019). Here, using the well-studied kinase BIN2, we demonstrate the application of TurboID-mediated PL-MS in dissecting signaling networks.

PL has not been used extensively in plants. Recent studies using BioID has achieved limited success in identification of interactors of transcription factor in rice protoplasts (Lin et al., 2017), and of pathogen effectors in *Nicotiana benthamiania* and Arabidopsis (Conlan et al., 2018, Khan et al., 2018). In all these studies biotinylation by BioID requires a long time (24 hr) treatment with a high concentration of biotin (>1 mM) (Khan et al., 2018). Our results show that biotinylation of cellular proteins in Arabidopsis expressing BIN2-YFP-TbID can be detected within one hour of treatment with a low concentration of 5~50 μM biotin. The increased labeling efficiency of TurboID not only improves detection sensitivity but also improves time resolution, which is important for analyzing PPI dynamics and identifying proteins that turn over rapidly *in vivo*.

Our direct comparison between PL and IP from the same BIN2-YFP-TbID extract shows that PL with TurboID is about ten times more effective in pulling down BZR1 by streptavidin than by IP with anti-GFP antibody. It is worth noting that BIN2 binds to a 12-amino acid motif of BZR1, and this kinase-substrate interaction involves a direct docking mechanism and is likely more stable than many other kinase-substrate interactions (Peng et al., 2010). Bigger improvement by PL-MS over IP-MS would be expected for more transient interactors of BIN2.

The condition of biotin treatment is important for optimal sensitivity and specificity of PL-MS. The promiscuous biotin ligases is known to catalyze the formation of biotin-5′-AMP anhydride, which diffuses out of the active site to biotinylate proximal endogenous proteins within a radius of about 10 nm for BioID (Kim et al., 2014). TurboID appears to have similar labeling radius as BioID, despite much faster labeling kinetics (Branon et al., 2018). However, long time biotin treatment increases biotinylation of non-specific proteins, probably due to saturation of the proximal sites and increase of labeling radius (Branon et al., 2018). Long time labeling can not only decrease the spatial specificity but also interfere with normal cellular processes via excessive biotinylation of endogenous proteomes (Branon et al., 2018). It is therefore important to empirically optimize the *in vivo* labeling time window, and use the shortest labeling time that produces sufficient biotinylated material for analysis.

We showed that the amount of biotinylated protein increased with the increases of biotin concentration and time of treatment. MS analysis of biotinylated peptides after treatment with different concentrations of biotin showed saturation of biotinylation of a set of proteins at a low concentration of 5 μM biotin, whereas biotinylation of additional proteins increase quantitatively with increasing biotin concentration. Interestingly, the BIN2-proximal proteins identified using YFP-YFP-TbID control (Figure 5-source data 2) were mostly detected in samples treated with 5 μM biotin (109 proteins) (Figure 5-supplemental figure 2). Increasing the biotin concentrations to 50 and 500 μM each detected only three additional BIN2-proximal proteins, among the 78 and 216 additional biotinylated proteins detected at these conditions (Figure 5-supplemental figure 2). It is conceivable that biotinylation of a set of direct or stable interactors is saturated at low concentrations of biotin, whereas treatment with higher concentrations of biotin lead to cumulative biotinylation of additional proteins outside the initial labeling radius or proteins that interact with BIN2 dynamically.

In addition to affecting sensitivity and specificity, use of high concentrations of biotin can greatly reduce pulldown efficiency if not removed from the extracts. Apparently, the plant tissues can carry over significant amount of free biotin into the extracts, which compete for binding to streptavidin. Removing free biotin from the extract, e.g. by using a desalting column or protein precipitation, can greatly increase the pulldown efficiency. We also found that streptavidin pulldown from protein extracts under mild buffer conditions identified fewer biotinylated proteins than pulldown of tryptic peptides under a more stringent condition, likely because of higher background binding of proteins than peptides. While purification at protein level may have higher MS detection sensitivity than peptide purification because each protein generates multiple peptide fragment ions, purification at peptide level would reduce sample complexity for MS as well as the level of non-specific noise. These results suggest that these two purification methods are complementary, and combining the two purification methods, which can be further optimized, will improve data coverage.

The specificity of PL in detecting PPIs needs evaluation. Direct comparison between known interactors and non-interactors within the GSK3 and PP2AB’ family members show that TurboID has a high level of specificity that distinguishes members of these protein families. Nevertheless, as non-specific cellular proteins have chances to enter the labeling radius and become biotinylated, quantitative comparison to a proper negative control is crucial for distinguishing specific interactors from non-specific background in PL-MS, like for IP-MS. We used a YFP-YFP-TurboID as our negative control for BIN2-YFP-TbID, and used the YFP tag to verify that the control and experimental constructs had similar subcellular localization. Among the 55 BIN2-proximal proteins identified in the protein AP and mSIL-MS experiment, 25 proteins were detected in the peptide AP-MS experiments, and 24 of them showed higher biotinylation by BIN2-TurboID in both experiments, indicating a high consistency of PL-MS in identifying proximal proteins. However, the partial overlap between the data of two experiments indicate that there are still false negative results due to incomplete MS coverage. Further, some BIN2 interactors may happen to be biotinylated also by the YFP-TurboID control, resulting in a fold enrichment below the cutoff threshold.

Among the 310 BIN2-proximal proteins we identified are eight BIN2-interactors that were previously reported. These include BIN2’s substrates BZR1, BZR2/BES1 (He et al., 2002), ARF2 (Vert et al., 2008), YDA (Kim et al., 2012), and its upstream regulators BSK3 (Tang et al., 2008, Sreeramulu et al., 2013), BSL2, BSL3 (Kim et al., 2009), and OCTOPUS (Anne et al., 2015). Many of the previously reported BIN2 interactors were not identified in our PL-MS experiments. These include BIN2’s E3 ligase KIB1 (Zhu et al., 2017), which might be turned over together with ubiquitinated BIN2. The second class of undetected BIN2 interactors are cell-type specific. These include the xylem differentiation-regulating receptor kinase TDR, stomatal lineage cell-specific SPCH, BASL and POLAR (Gudesblat et al., 2012, Kondo et al., 2014, Cho et al., 2014, Houbaert et al., 2018). The third class of reported BIN2 interactors that were undetected in our PL-MS experiments include AIF1, CESTA, MYBL2, and Tubulins; the discrepancies will need to be resolved in future studies.

The large number of BIN2-proximal proteins identified in our study suggests very broad BIN2 signaling networks. Some of the new BIN2-proximal proteins potentially mediate known functions of the BR pathway. For example, BIN2 interaction with the auxin transporter PIN3 may explain the observation that BR modulates polar auxin transport and gravitropic responses (Bao et al., 2004, Kim et al., 2000). BR has also been reported to play a role in phototropism (Whippo and Hangarter, 2005), and our PL-MS data indicate that BIN2 interacts with proteins involved in phototropism: Phototropin1 (PHOT1), NONPHOTOTROPIC HYPOCOTYL3 (NPH3), and a NPH3 family protein. We also identified six receptor kinases, including HERK1 and FER, which have been reported to play a role in BR-mediated growth responses (Guo et al., 2009). The large number of BIN2-proximal proteins supports the developing idea that BIN2 may play broader roles in additional signaling pathways beyond BR signaling. Functional and phospho-proteomic studies of these BIN2-proximal proteins will further advance toward a complete understanding of the BR/BIN2 signaling network. Alternatively, similar PL-MS analysis of GSK3s in other plant species will provide orthogonal information about conservation and thus functional constrains of the network (Levy et al., 2009).

Our study demonstrates that TurboID is a powerful tool for dissecting signaling networks. TurboID is also powerful for analyzing the proteomes of specific cell types and even subcellular compartments in plants (Mair, 2019)(accompanying manuscript). Combining these two applications, by expressing the TurboID fusion with signaling proteins using cell type-specific promoters, will not only improve the sensitivity and specificity of the analysis but also illustrate signaling network in specific developmental context.

## Materials and methods

### Plant Materials and Growth Condition

*Nicotiana benthamiana* seeds were planted on soil and grown for 4-5 weeks in the green house. *YFP-YFP-TbID* and *BIN2-YFP-TbID* were overexpressed in Arabidopsis Col-0 ecotype. Arabidopsis seedlings were grown on 1/2 Murashige and Skoog medium (MS; PhytoTechnology Laboratories, Shawnee Mission, KS) containing 1% (w/v) sucrose and 0.8% phytoagar (Caisson Laboratories, East Smithfield, UT).

### Plasmids

To generate a Gateway-compatible 35S-YFP-TbID vector, PCR fragments obtained from TurboID-containing plasmid (V5-TbID-NES_pCDNA3, Addgene) and pEarleyGate101 vector were assembled by overlapping ends using Gibson assembly master mix (NEB, Ipswich, MA). TurboID was amplified with primers FP1 (5’-ATCCACCGGATCTAGAGGCAAGCCCATCCCCAAC-3’) and RP1 (5’-AACATCGTATGGGTAAGGCAGCTGCAGCTTTTCGG-3’) while the pEarleyGate101 vector was amplified with primers FP2 (5’-TACCCATACGATGTTCCAGATTACGCTTAATTAA-3’) and RP2 (5’-CTTGCCTCTAGATCCGGTGGATCCC-3’). The coding sequences of YFP and BIN2 cloned in pENTR/SD/D-TOPO were subcloned into a Gateway-compatible 35S-YFP-TurboID by LR reaction (Invitrogen, Carlsbad, CA).

### *Nicotiana benthamiana* Infiltration

Agrobacterium was inoculated into 5 mL of LB medium and grown for 16 hours at 28 °C. Cultured cells harvested from 1 mL aliquot were resuspended with 2 mL of the induction media (10 mM MES pH 5.6, 10 mM MgCl_2_, and 150 μM Acetosyringone), mixed according to the combination of plasmids, and incubated for 1 h at room temperature. Cells were infiltrated into abaxial leaves of *Nicotiana benthamiana* using 1 mL syringe. After 36~40 hours, leaves were harvested and kept at −80 °C until use.

### Protein Extraction and Immunoblot Analysis

Plant tissues were ground with liquid nitrogen and sample powder was resuspended with two volumes (2 ml per gram tissue) of extraction buffer (20 mM HEPES, pH 7.5, 40 mM KCl, 1 mM EDTA, 1% Triton X-100, 0.2 mM PMSF, and 1X protease inhibitor cocktail). The protein mixtures were centrifuged at 4,000 rpm for 5 min. Then, resulting supernatant was centrifuged at 20,000 g for 15 min. The supernatants were incubated with Streptavidin-agarose (S1638, Sigma, Saint Louis, MO) or an anti-Myc-tag Nanobody coupled to agarose (Myc-Trap, Chromotek, Hauppauge, NY) for 1 hour at 4 °C. Then, beads were washed with an extraction buffer containing 0.1% Triton X-100 and eluted with 2X SDS sample buffer (24 mM Tris-Cl, pH 6.8, 10% glycerol, 0.8% SDS, 2% 2-mercaptoethanol) containing 0.4 M urea. YFP-TbID and BZR1-MH were detected by monoclonal anti-GFP (HT801, Transgen Biotech, Beijing, China) and monoclonal anti-Myc antibodies (9B11, Cell Signaling Technology, Danvers, MA), respectively. Biotinylated proteins were detected with streptavidin-HRP (21124, Thermo Scientific, Rockford, IL).

### Removing free biotin and affinity purification of biotinylated proteins from total extract of BIN2-YFP-TbID

To obtain protein extracts, BIN2-YFP-TbID seedlings treated with 50 μM biotin for 3 hours were ground with liquid nitrogen. One gram of the tissue powder was resuspended with 1.5 mL IP buffer (50 mM Tris, pH 7.5, 50 mM NaF, 300 mM sucrose, 1% Triton X-100) with 1x protease inhibitor cocktail (Pierce, Rockford, IL) and centrifuged at 1,500 rpm for 5 min. Resulting supernatant was centrifuged at 12,000 rpm for 10 min and each 1 mL aliquot of the supernatant was transferred to two tubes. For desalting, one of the 1 mL supernatant was desalted by PD-10 desalting columns (GE Healthcare, Pittsburgh, PA) according to manufacturer’s instructions and eluted with 3 mL IP buffer. To the other supernatant sample, 2 mL IP buffer was added to make the same volume with desalted sample. The protein samples were incubated with 30 μL Dynabead C1 Streptavidin beads (Thermo Fisher Scientific, Waltham, MA) at 4 °C for 3 hours. The beads were subsequently washed 3 times with washing buffer (50 mM Tris, pH 7.5, 100 mM NaCl, 50 mM NaF, 0.1% Triton X-100). Biotinylated proteins were eluted by boiling the bead in 50 μL 2× SDS sample buffer (100 mM Tris, pH 6.8, 4% sodium dodecyl sulfate, 20% glycerol and 100 mM dithiothreitol) and separated by a SDS-PAGE gel (Biorad, Hercules, CA). Biotinylated protein was detected with streptavidin-HRP (21124, Thermo Scientific, Rockford, IL).

### Confocal Microscopy

*BIN2-YFP-TbID* and *YFP-YFP-TbID* seedlings were grown on MS agar medium for 8 days. YFP fluorescence of root segments was visualized with an SP8 confocal microscope (Leica Microsystems, Heerbrugg, Germany).

### Metabolic Stable Isotope Labeling and Affinity Purification of Biotinylated Proteins

Metabolic stable isotope labeling (SIL) of Arabidopsis seedlings was performed as follows. Transgenic *BIN2-YFP-TbID* and *YFP-TbID* seedlings were grown on ^14^N MS medium (1/2 MS without nitrogen source [PhytoTechnology Laboratories], NH_4_NO_3_ [0.5 g/L, Sigma], KNO3 [0.5 g/L, Sigma], pH 5.7) or ^15^N MS medium (1/2 MS without nitrogen source [PhytoTechnology Laboratories], ^15^NH4^15^NO_3_ [0.5 g/L, NLM-390-1, Cambridge Isotope Laboratory], K^15^NO_3_ [0.5 g/L, NLM-765-1, Cambridge Isotope Laboratory], pH 5.7) for 16 days under continuous light in a growth chamber at 22 °C. ^14^N- and ^15^N-labeled seedlings were harvested and syringe infiltrated with 50 μM biotin solution. After infiltration, seedlings were transferred to 50 mL conical tubes and further incubated with 50 μM biotin for 3 hours in the growth chamber. Proteins from 5 g tissue powder ground in liquid nitrogen were extracted with 5 mL IP buffer (50 mM Tris, pH 7.5, 50 mM NaF, 300 mM sucrose, 1% Triton X-100) containing 1x protease inhibitor cocktail (Pierce, Rockford, IL) and a phosphatase inhibitor (PhosSTOP, Roche Applied Science, Penzberg, Germany). Protein extracts were centrifuged at 1,500 rpm for 5 min and the supernatant was re-centrifuged at 12,000 rpm for 10 min. To remove free biotin, the resulting supernatant was desalted by PD-10 desalting columns (GE Healthcare, Pittsburgh, PA) according to manufacturer’s instructions. Elute was incubated with 100 μL Dynabead M-280 Streptavidin beads (Thermo Fisher Scientific, Waltham, MA) at 4 °C for 8 hours. The beads were subsequently washed 3 times with washing buffer (50 mM Tris, pH 7.5, 100 mM NaCl, 50 mM NaF, 0.1% Triton X-100). The beads from ^14^N-labeled *BIN2-TbID* were transferred to new low protein binding tube (Thermo Fisher Scientific) and mixed with that of ^15^N-labeled *YFP-TbID*. The mixed beads were washed once with the washing buffer and biotinylated proteins were eluted by 30 μL 2× SDS sample buffer and separated by SDS-PAGE (Biorad, Hercules, CA).

### In-gel Trypsin Digestion

Protein bands were stained by Colloidal Blue staining (Invitrogen). Four segments of the protein bands were excised into 1 mm blocks. Protein samples were reduced with 10 mM dithiothreitol for 1 hour at 56 °C. The samples were alkylated with 50 mM iodoacetamide for 45 min in the dark at room temperature. Trypsin digestion was performed by adding 0.25 μg of trypsin to the sample and the digestion was completed overnight at 37 °C. Digested peptides were extracted from the gel by peptide extraction buffer (50% acetonitrile, 0.1% formic acid) and dried in a vacuum concentrator (Thermo Fisher Scientific). Peptides were resuspended in 0.1% formic acid and desalted using C18 ZipTips (Millipore, Burlington, MA), and analyzed by LC-MS/MS.

### LC-MS/MS Analysis of SIL samples

The peptides were analyzed on a Q-Exactive HF hybrid quadrupole-Orbitrap mass spectrometer (Thermo Fisher) equipped with an Easy LC 1200 UPLC liquid chromatography system (Thermo Fisher). Peptides were separated using analytical column ES802 (Thermo Fisher). The flow rate was 300 nL/min and a 120-min gradient was used. Peptides were eluted by a gradient from 3 to 28% solvent B (80% acetonitrile/0.1 formic acid) over 100 mins and from 28 to 44% solvent B over 20 mins, followed by short wash at 90% solvent B. Precursor scan was from mass-to-charge ratio (m/z) 375 to 1600 and top 20 most intense multiply charged precursor were selected for fragmentation. Peptides were fragmented with higher-energy collision dissociation (HCD) with normalized collision energy (NCE) 27. MS/MS data was converted to peaklist using an in-house script PAVA, and data was searched using Batch-Tag of Protein Prospector against the TAIR database *Arabidopsis thaliana* (https://www.arabidopsis.org/), concatenated with sequence randomized versions of each protein (a total of 35386 entries). A precursor mass tolerance of 10 ppm and a fragment mass error tolerance of 20 ppm were allowed. Carbamidomethylcysteine was searched as a constant modification. Variable modifications include protein N-terminal acetylation, peptide N-terminal Gln conversion to pyroglutamate, and Met oxidation.

Median intensity ratio between light and heavy peaks (Med L/H I) was used for protein level quantitation. Average of Med L/H I value from ^14^N and ^15^N searches was calculated. Proteins with the BIN2-TbID/YFP-TbID ratio ≥2 in both repeat experiments were considered BIN2-proximity labelled proteins. Scatter plot of log2 fold change values of BIN2-enriched proteins were generated by SigmaPlot.

### Streptavidin purification of biotinylated peptides

Protein extraction and digestion was performed as previously described (Hsu et al., 2018). Plants were lysed in lysis buffer (6M guanidine hydrochloride in 100 mM Tris-HCl at pH 8.5) with EDTA-free protease and phosphatase inhibitor cocktails. Disulfide bond on proteins was reduced and alkylated with 10 mM Tris (2-carboxyethyl) phosphine hydrochloride and 40 mM 2-chloroacetamide at 95 °C for 5 min. Protein lysate was precipitated using methanol-chloroform precipitation method. Precipitated protein pallets were suspended and in digestion buffer (12 mM sodium deoxycholate and 12 mM sodium N-Lauroyl sarcosine in 100 mM Tris-HCl at pH 8.5) and then were 5-fold diluted with 50 mM triethylammonium bicarbonate buffer. Protein amount was quantified using BCA assay (Thermo Fisher Scientific). Five mg of proteins were then digested with Lys-C (Wako) in a 1:50 (v/w) enzyme-to-protein ratio for 3 hours at 37 °C, and trypsin (Sigma) was added to a final 1:100 (w/w) enzyme-to-protein ratio overnight. The detergents were separated from digested peptides by acidifying the solution using 10% trifluoroacetic acid and then centrifuged at 16,000 g for 20 min. The digests were then desalted using a 100 mg SEP-PAK C18 cartridge (Waters).

Biotinylated peptides were enriched using Dynabeads M-280 Streptavidin (Thermo Fisher Scientific). The digested peptides were resuspened with 1 ml of PBS buffer, and then 100 μl of Dynabeads were added to the sample and incubated for 1 hour at room temperature. The solution was removed using a magnetic rack, and Dynabeads were washed with 1 ml of PBS buffer for 5 min twice. The beads then further incubated with 1 ml of washing buffer 1 (PBS with 5% acetonitrile) for 5 min twice, and then were incubated with 1 ml of washing buffer 2 (dd¾O with 5% acetonitrile) for 5 min twice. Finally, the biotinylated peptides were eluted with 200 μl of elution buffer (80% acetonitrile with 0.2% trifluoroacetic acid and 0.1% formic acid) for 5 min five times. The eluates were combined and dried using a SpeedVac, and then were desalted using a C18 StageTip.

### Streptavidin purification of biotinylated peptides

Protein extraction and digestion was performed as previously described (Hsu et al., 2018). Plants were lysed in lysis buffer (6M guanidine hydrochloride in 100 mM Tris-HCl (pH 8.5)) with EDTA-free protease and phosphatase inhibitor cocktails. Disulfide bond on proteins was reduced and alkylated with 10 mM Tris(2-carboxyethyl)phosphine hydrochloride and 40 mM 2-chloroacetamide at 95 °C for 5 min. Protein lysate was precipitated using methanol-chloroform precipitation method. Precipitated protein pallets were suspended in digestion buffer (12 mM Sodium deoxycholate and 12 mM Sodium lauroyl sarcosinate in 100 mM Tris-HCl, pH 8.5) and then were 5-fold diluted with 50 mM triethylammonium bicarbonate buffer. Protein amount was quantified using BCA assay (Thermo Fisher Scientific). Five mg of proteins were then digested with Lys-C (FUJIFILM Wako) in a 1:50 (v/w) enzyme-to-protein ratio for 3 hours at 37 °C, and trypsin (Sigma-Aldrich) was added to a final 1:100 (w/w) enzyme-to-protein ratio overnight. The detergents were separated from digested peptides by acidifying the solution using 10% trifluoroacetic acid (TFA) and then centrifuged at 16,000 g for 20 min. The digests were then desalted using a 100 mg SEP-PAK C18 cartridge (Waters).

The digested peptides were resuspended with 1 ml of PBS buffer, and then 100 μl of the Dynabeads M-280 Streptavidin (Thermo Fisher Scientific) beads were added to the sample and incubated for 1 hour at room temperature. The solution was removed using a magnetic rack, and the Dynabeads were washed with 1 ml of PBS buffer for 5 min twice. The beads then further incubated with 1 ml of washing buffer 1 (PBS with 5% acetonitrile (ACN)) for 5 min twice, and then were incubated with 1 ml of washing buffer 2 (ddH_2_O with 5% ACN) for 5 min twice. Finally, the biotinylated peptides were eluted with 200 μl of elution buffer (80% ACN with 0.2% TFA and 0.1% formic acid) for 5 min five times. The eluates were combined and dried using a SpeedVac, and then were desalted using a C18 StageTip(Rappsilber et al., 2007).

### LC-MS/MS Analysis for biotinylated peptide detection

The peptides were dissolved in 5 μL of 0.3% formic acid with 3% acetonitrile and injected into an Easy-nLC 1200 (Thermo Fisher Scientific). Peptides were separated on a 25 cm Easy-Spray column (75 μm ID) containing C18 resin (1.9 μm) with a column heater set at 40 °C. The mobile phase buffer consisted of 0.1% formic acid in ultra-pure water (buffer A) with an eluting buffer of 0.1% formic acid in 80% acetonitrile (buffer B) run over a linear 65 min (method comparisons) gradient of 5%-28% buffer B at a flow rate of 300 nL/min. The Easy-nLC 1200 was coupled online with a Q-Exactive HF Orbitrap mass spectrometer (Thermo Fisher Scientific). The mass spectrometer was operated in the data-dependent mode in which a full MS scan (from m/z 375-1600 with the resolution of 60,000 at m/z 400) was followed by the 10 most intense ions being subjected to higher-energy collision dissociation (HCD) fragmentation. HCD fragmentation was performed and acquired in the Orbitrap (normalized collision energy (NCE) 27%, AGC 3e4, max injection time 120 ms, isolation window 1.5 m/z, and dynamic exclusion 25 s).

### Data Processing and Analysis for biotinylated peptide detection

The raw files were searched against a TAIR10 database (35,386 entries) from TAIR (https://www.arabidopsis.org/) using MaxQuant software (version 1.6.2.10) (Cox and Mann, 2008) with a 1% FDR cutoff at the protein, peptide, and modification levels. The first peptide precursor mass tolerance was set at 20 ppm, and MS/MS match tolerance was set at 20 ppm. Search criteria included a static carbamidomethylation of cysteines and variable modifications of (1) oxidation on methionine residues, (2) acetylation at the N-terminus of proteins, and (3) biotinylation (+ 226.078 Da) on N-terminus of proteins or lysine residues. The match between runs function was enabled with a match time window of 0.7 min. The minimum peptide length was seven amino acids, and a minimum Andromeda score was set at 40 for modified peptides. All data were analyzed using the Perseus software (version 1.6.2.1) (Tyanova et al., 2016) and Microsoft Excel. The number of unique biotinylated peptides identified from each samples was calculated using Microsoft Excel after removal of the redundant biotinylated sequences in the MaxQuant output modificationSpecificPeptide file. The intensities of biotinylated peptides were log2 transformed, and the quantifiable biotinylated peptides were selected from the identification of all triplicate replicates from BIN2-tbID samples but not in any of YFP-tbID samples.

### Data Availability

The mass spectrometry proteomics data have been deposited to the ProteomeXchange Consortium (Vizcaino et al., 2014) with the dataset identified PXD013768. (username, password: KqYUvh9L).

## Acknowledgements

This research was supported by a grant from NIH (R01GM066258 to Z-Y. W.) and by Carnegie endowment fund to the Carnegie mass spectrometry facility. This research was also supported by the Basic Science Research Program through the National Research Foundation of Korea (NRF) funded by the Ministry of Science, ICT, and Future Planning (NRF-2017R1A2B4004274 to T.W.K.).

## Author contributions

Tae-Wuk Kim, Conceptualization, Validation, Investigation, Resources, Writing—original draft, Writing—review and editing, Visualization, Supervision, Project administration; Chan Ho Park, Conceptualization, Validation, Formal analysis, Investigation, Resources, Data curation, Writing—original draft, Writing—review and editing, Visualization; Chuan-Chih Hsu, Validation, Investigation, Data curation, Visualization; Jia-Ying Zhu, Resources; Yuchun Hsiao, Resources; Tess Branon, Resources; Shouling Xu, Formal analysis; Alice Y Ting, Conceptualization, Methodology, Resources; Zhi-Yong Wang, Conceptualization, Writing— original draft, Writing—review and editing, Supervision, Project administration, Funding acquisition.

---

**Figure 2-Figure supplement 1.**
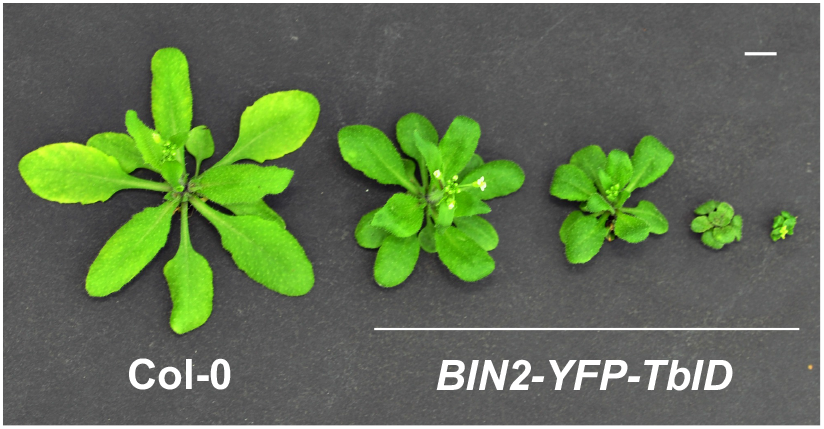
Growth phenotypes of 4-week-old wild-type and *BIN2-YFP-TblD* Arabidopsis plants. Bar = 1 cm.

**Figure 4-Figure supplement 1.**
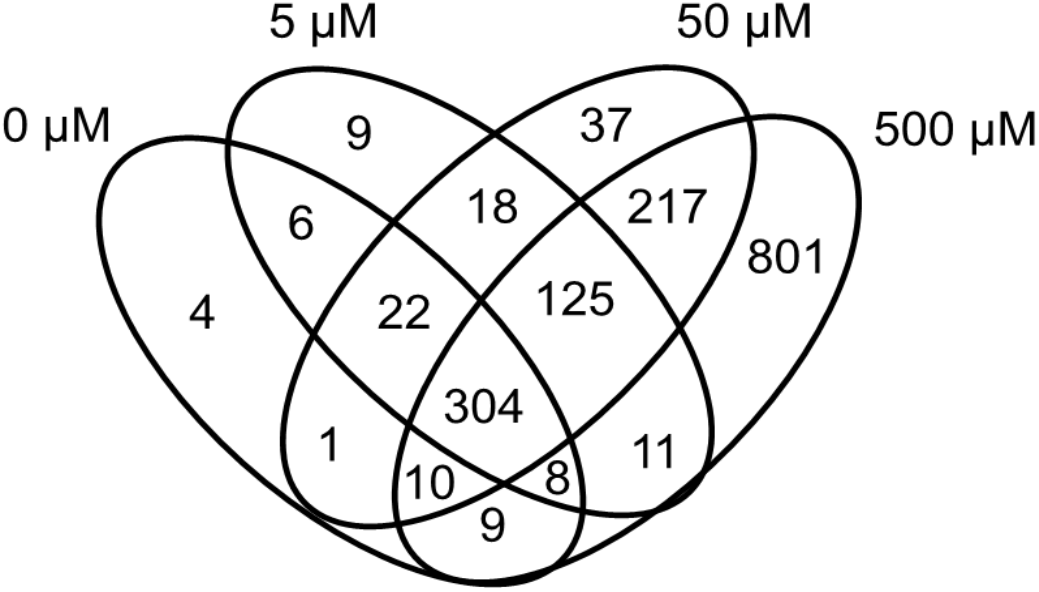
The overlaps of identified biotinylated peptides between samples treated for 1 hour with 0, 5, 50, and 500 μM biotin.

**Figure 5-Figure supplement 1.**
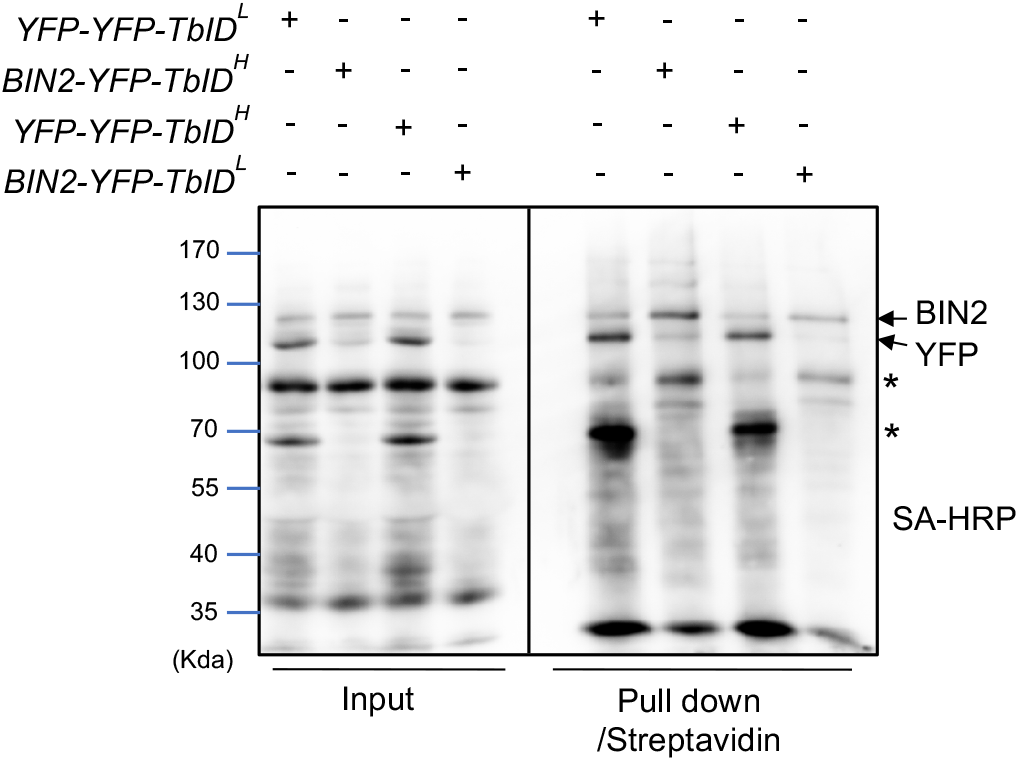
Gel blot analysis of mSIL-MS samples. Biotinylated proteins from ^14^N-labeled (^L^) and ^15^N-labeled (^H^) *BIN2-YFP-TbID* and *YFP-YFP-TbID* were pulled down by Streptavidin-coated magnetic beads. Input (1:700) and eluted pull down (1:40) were separated by SDS-PAGE. Total biotinylated proteins were probed by Streptavidin-HRP (SA-HRP). Arrows indicate full length and self-biotinylated BIN2-YFP-TbID (BIN2) and YFP-YFP-TbID (YFP) proteins. Asterisks indicate truncated proteins.

**Figure 5-Figure supplement 2.**
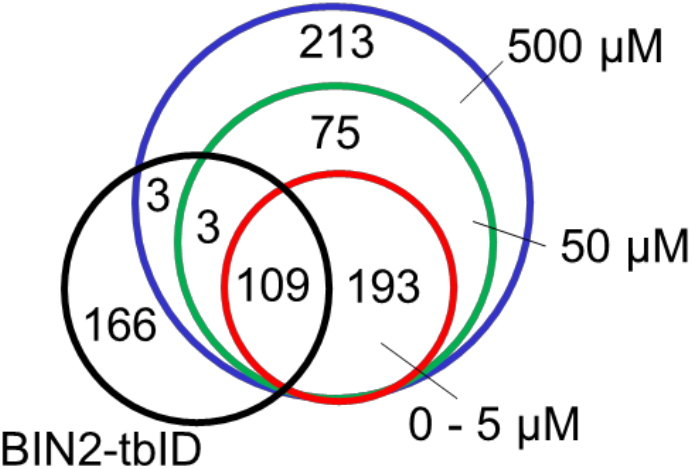
The overlap of biotinylated proteins identified in samples treated with different concentrations of biotin (0~5, 50, and 500 μM) with the BIN2-proximal proteins (BIN2-TbID) identified by comparing BIN2-TbID with YFP-TbID control (Figure 5-Source data 2).

